# Metabolic labeling of RNA using multiple ribonucleoside analogs enables simultaneous evaluation of transcription and degradation rates

**DOI:** 10.1101/2020.03.06.980250

**Authors:** Kentaro Kawata, Hiroyasu Wakida, Toshimichi Yamada, Kenzui Taniue, Han Han, Masahide Seki, Yutaka Suzuki, Nobuyoshi Akimitsu

## Abstract

Gene expression is one of the factors determining cellular conditions, and its level is determined by the balance between transcription and RNA degradation. However, it has been not well elucidated how transcription and degradation are related to each other. To elucidate the relationship of them, methods enabling simultaneous measurement of transcription and degradation are required. Here, we report the development of “Dyrec-seq” to evaluate transcription and RNA degradation rates simultaneously, by using 4-thiouridine and 5-bromouridine. Using Dyrec-seq, we quantified the transcription and degradation rates of 4,702 genes in HeLa cells. Functional enrichment analysis showed that the transcription and degradation rates of genes are actually determined by the genes’ biological functions. Comparison of theoretical and experimental analysis results revealed that the amount of RNA was determined by the ratio of transcription to degradation rates, while the rapidity of responses to external stimuli was determined only by the degradation rate. This study emphasizes that degradation as well as transcription is important in determining the behavior of RNA.

## Introduction

Gene expression is one of the most fundamental regulatory processes determining cellular conditions by regulating the level of proteins. RNAs are generated as initial products of gene expression, the levels of which are determined through complex processes including transcription and degradation (Rabani *et al*., 2011, 2014; De Pretis *et al*., 2015; Maekawa *et al*., 2015; McManus, Cheng and Vogel, 2015; Eser *et al*., 2016; Baptista and Dölken, 2018; Kiefer, Schofield and Simon, 2018; Duffy, Schofield and Simon, 2019; Schmid, Tudek and Jensen, 2019). The rate of transcription of a gene is regulated by complex interactions of regulatory *cis*-acting elements including promoter and enhancer regions in the genome, and *trans*-acting elements such as transcription factors (TFs) and epigenetic regulators. In contrast, the rate of degradation of RNA is regulated by several RNA degradation complexes consisting of RNA-binding proteins (RBPs) and microRNAs (miRNAs). To obtain a comprehensive overview of the regulation of gene expression, simultaneous measurement of different regulatory steps, such as transcription and RNA degradation, is required. A range of procedures are available to measure the rate of either transcription or degradation of each RNA at the genome-wide level (Tani *et al*., 2012; Imamachi *et al*., 2014; Schwalb *et al*., 2016; Herzog *et al*., 2017; Baptista and Dölken, 2018; Kiefer, Schofield and Simon, 2018; Matsushima *et al*., 2018; Schofield *et al*., 2018; Duffy, Schofield and Simon, 2019). For example, SLAM-seq enables the transcription rates of RNAs to be measured using *in situ* labeling of RNAs with 4-thiouridine (4sU) (Herzog *et al*., 2017). The 4sU is alkylated *in vitro* after isolation of total RNA from 4sU-labeled cells, followed by massive sequencing analysis. Since a guanine (G) instead of an adenine (A) pairs with alkylated 4sU during the reverse-transcription reaction in library preparation for massive sequencing, a thymine (T) for the nascent RNA in sequencing data is converted to a cytosine (C) (T C conversion). Bioinformatic detection of T > C conversion enables us to distinguish the nascent RNA from preexisting RNAs. By measuring T > C conversion in RNAs at sequential time points after 4sU labeling, the transcription rates of each RNA can be determined. BRIC-seq enables measurement of the degradation rates of the RNAs using *in situ* labeling of RNA with 5′-bromouridine (BrU) (Tani *et al*., 2012; Imamachi *et al*., 2014). The BrU-labeled RNAs chronologically isolated from cells pre-labeled with BrU are immunoprecipitated, and then the immunoprecipitated RNAs are chased by massive sequencing to estimate the degradation rates of the RNAs. However, methods enabling simultaneous measurement of transcription and degradation by using multiple ribonucleoside analogs are not currently available.

In this study, we first simultaneously measured actual transcription and degradation rates in the human cervical cancer HeLa cell line at the genome-wide level, by developing “sequencing for RNA <u>dy</u>namics <u>rec</u>ording” (Dyrec-seq), combining SLAM-seq and BRIC-seq. Using Dyrec-seq, we determined the transcription and degradation rates of 4,702 genes. Functional enrichment analysis indicated that the transcription and degradation rates of genes are actually determined by the genes’ biological functions. How does the combination of the transcription and degradation rates affect gene expression? Theoretical analysis using a mathematical model clarified that the expression level of each RNA in a steady state is determined by ratio of the transcription rate to the degradation rate. Comparison of actually measured expression levels to the ratios of the transcription and degradation rates proved this theoretical prediction experimentally. Moreover, approaches of simulating the differential expression of genes and experimental validation using a published dataset indicated that the rapidity of responses to extracellular stimulation can be determined by the degradation rate, but not by the transcription rate. Overall, Dyrec-seq clarified that the transcription and degradation rates of individual genes cooperatively regulate the behavior of gene expression, such as the expression level and rapidity of differential expression in response to environmental stimuli.

## Results

### Sequential labeling of endogenous RNAs with BrU and 4sU

To quantify transcription and degradation rates simultaneously, we labeled HeLa cells with BrU followed by 4sU (Fig. 1A). The HeLa cells were pre-cultured in BrU-containing medium for 12 h, and then RNAs were isolated at 0, 15, 30, 45, 60, 120, 240, 480, and 720 min after the BrU-containing medium had been changed to 4sU-containing medium without BrU (Total-RNAs). After 4sU was added and BrU was removed from the medium by changing the medium (time: 0 min), the abundances of BrU-labeled and 4sU-labeled RNAs decreased and increased, respectively, over time. The isolated RNAs were divided into two samples. One was immunoprecipitated with anti-BrdU antibody to isolate BrU-labeled RNAs (IP-RNAs) and the other was treated with iodoacetamide (IAA) to alkylate the 4sU residues in newly transcribed RNAs (Alkyl-RNAs) (Fig. 1A and B). The 3′ ends of RNAs within the Alkyl-RNAs and IP-RNAs were sequenced using QuantSeq (Moll *et al*., 2014), which enabled quantification of the mRNA expression by sequencing the sequences close to the 3′ end of polyadenylated RNA (Fig. 1C).

**Figure 1.**
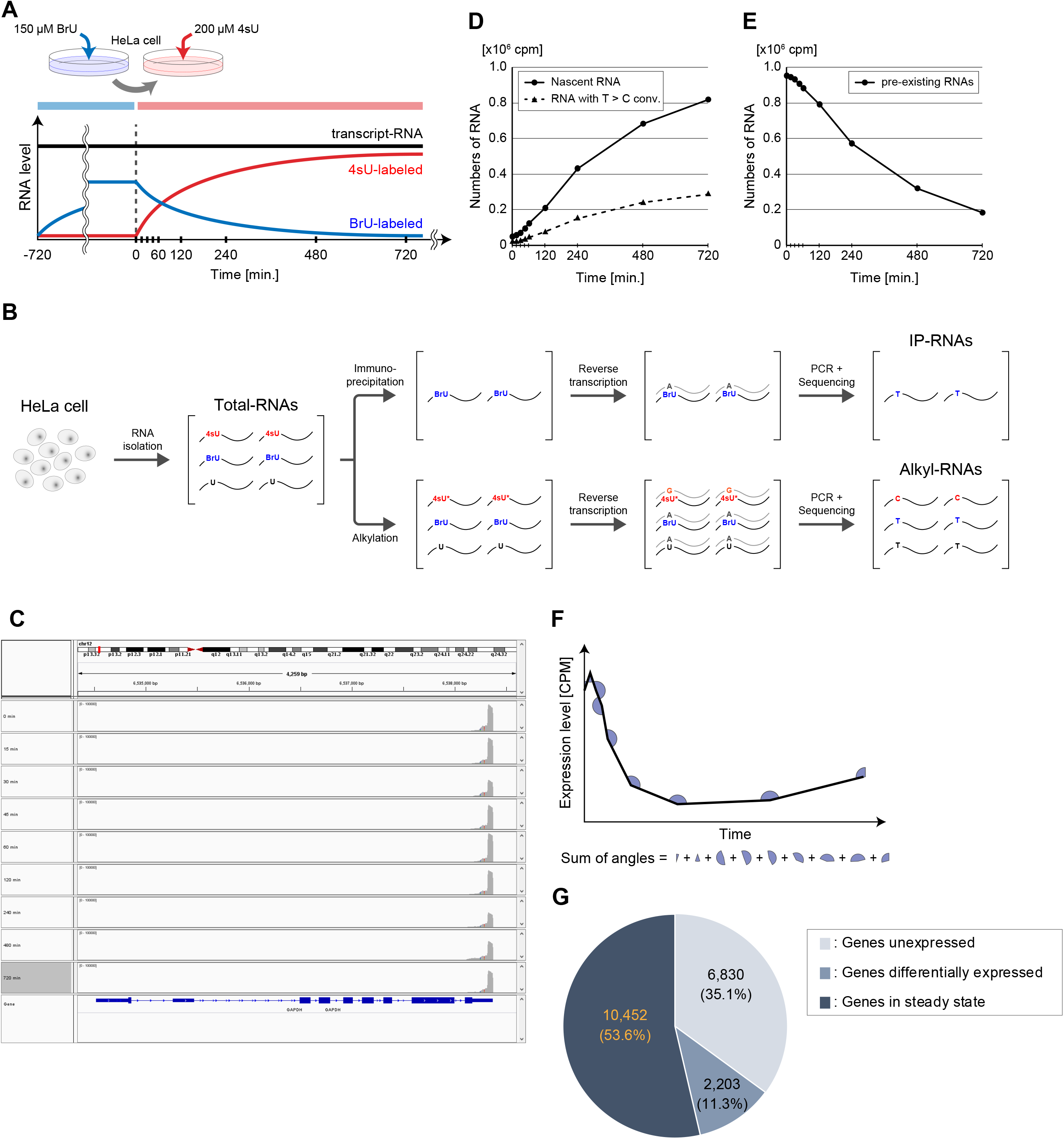
Simultaneous labeling of intracellular RNAs with BrU and 4sU. (**A**) Schematic illustration of the labeling procedure. HeLa cells were pre-cultured in medium containing 150 μM BrU and then the medium was changed to one containing 200 μM 4sU. Black, blue, and red lines indicate kinetics of expression of RNAs, BrU-labeled, and 4sU-labeled RNAs, respectively. (**B**) Preparation and quantification of BrU- and 4sU-labeled RNAs. Total RNAs were isolated and purified from labeled cells in time series, and then divided into two samples. One was immunoprecipitated using anti-BrdU antibody, and the cDNA was reverse-transcribed to be provided to QuantSeq (IP-RNAs). The other was alkylated using IAA and the cDNA was reverse-transcribed to be provided to QuantSeq (Alkyl-RNAs). Since alkylated 4sU pairs with guanine (G) instead of adenine (A) during reverse transcription, the 4sU-labeled RNAs were identified as those including T > C conversions. (**C**) A representative example of the alignment of reads sequenced using QuantSeq. The 3′UTRs of IP-RNAs and Alkyl-RNAs were sequenced using QuantSeq, poly(A)-dependent sequencing. (**D**) Estimation of nascent RNAs from sequencing of Alkyl-RNAs. The proportion of nascent RNAs incorporating 4sU was calculated by fitting a time series of RNAs including T > C conversions to a logarithmic curve (see STAR Methods). The amounts of nascent RNAs at individual time points were estimated by divided the RNAs including T > C conversions by the proportion of nascent RNAs incorporating 4sU. Solid and dashed lines indicate the time series of estimated nascent RNAs and RNAs with T > C conversions, respectively. (**E**) Estimation of pre-existing RNAs from the sequencing of IP-RNAs. Since the total amount of intracellular RNAs can be assumed to be constant during culture, the amount of pre-existing RNAs at each time point was estimated by subtracting the amount of estimated nascent RNAs from a constant (1 million). (**F**) Definition of the trend of gene expression. To remove the genes differentially expressed after 4sU labeling, a filter testing constant trends of expression was designed (see STAR Methods). The sum of the angles was defined as the sum of the angles formed by the lines connecting certain time points and neighboring ones, within a time series of total read numbers for each gene calculated from Alkyl-RNAs. (**G**) Distribution of gene expression state. Among the total of 19,485 annotated genes, 35.1%, 11.3%, and 53.6% were identified as those not expressed, differentially expressed, and in a steady state, respectively.

### Quantification of nascent and pre-existing RNAs

Subsequently, we estimated the numbers of nascent and pre-existing RNAs at each time point based on the sequencing of the Alkyl-RNAs and IP-RNAs by QuantSeq. Since 4sU is incorporated into nascent RNAs at a constant rate, the numbers of nascent RNAs are in proportion to the numbers of reads containing T > C mutations. Since nascent RNA is synthesized at a constant rate and degraded at a rate dependent on its concentration per unit time, the number of nascent RNAs increases logarithmically. Fitting of this curve of logarithmic increase indicated that approximately 35% of the nascent RNAs are labeled with 4sU (see STAR Methods). Therefore, we estimated the number of nascent RNAs by correcting their count per million mapped reads (CPM) using the labeling efficiency (Fig. 1D). Furthermore, we estimated the number of pre-existing RNAs using the total number of nascent RNAs at each time point. Since the total number of intracellular RNAs can be considered as constant and independent of time, the sum of the number of nascent and pre-existing RNAs should be constant. Thus, we estimated the number of pre-existing RNAs at each time point by subtracting the number of nascent RNAs at each time point from 1 million (Fig. 1E).

### Extraction of RNAs in a steady state

In this study, we assumed that the expression of genes was in a steady state during labeling with modified ribonucleosides. Although the cells were not treated with any specific stimuli, differential expression of some of the genes might actually have been attributable to the medium change or mechanical stress. Therefore, we extracted the genes in a steady state based on the statistical significance of their trends of expression (Fig. 1F) comparing their empirical distribution with that of a randomized gene set with preserved expression level (see STAR Methods). A statistical test indicated that, among 19,485 genes, 6,830 (35.1%) were not expressed at all time points, 2,203 (11.3%) were expressed differentially (*FDR* < 0.01), and 10,452 (53.6%) were in a steady state (*FDR* > 0.01) (Fig. 1G). We extracted these 10,452 genes in a steady state as subjects of further analysis.

### Simultaneous evaluation of transcription and degradation rates

Next, we estimated the transcription and degradation rates for individual genes simultaneously. The transcription rate of each gene is defined as the number of RNAs transcribed per unit time, and calculated based on the time series of the number of nascent RNAs estimated from the sequencing of the Alkyl-RNAs. The degradation rate of each RNA is defined as the ratio of the RNAs degraded per unit time calculated based on the time series of the number of pre-existing RNAs estimated from the sequencing of the IP-RNAs. Since the medium change removes BrU and adds 4sU, the numbers of reads derived from nascent and pre-existing RNAs increase and decrease in transcription rate- and degradation rate-dependent manners, respectively.

That is, the kinetics of the read numbers derived from nascent RNAs (*x*_*t*_) and pre-existing RNAs (*y*_*t*_) at each time point is as follows:

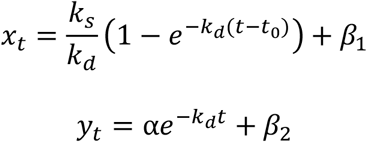

where *k*_*s*_, *k*_*d*_, *t*_0_, *α*, *β*_1_, and *β*_2_ are the transcription rate, degradation rate, time lag, scaling factor, basal value for nascent RNA-derived reads, and basal value for pre-existing RNA-derived reads, respectively (Fig. 2A–C). We estimated the transcription and degradation rates of individual genes at the genome-wide level by fitting the time series of the read numbers to these curves, and extracting well-fitted RNAs based on the statistical significance of correlation coefficients between the read numbers and estimated values (see STAR Methods). As a result, we obtained the transcription and degradation rates for 4,702 genes in HeLa cells (Table S1). The estimated transcription rates obeyed a log normal-like unimodal distribution with a median of 1.17×10^−1^ cpm·min^−1^ (Fig. 2D). The estimated degradation rates obeyed a log normal-like unimodal distribution with a median of 3.38×10^−3^ min^−1^, equivalent to a half-life of 205.0 min (Fig. 2E).

**Figure 2.**
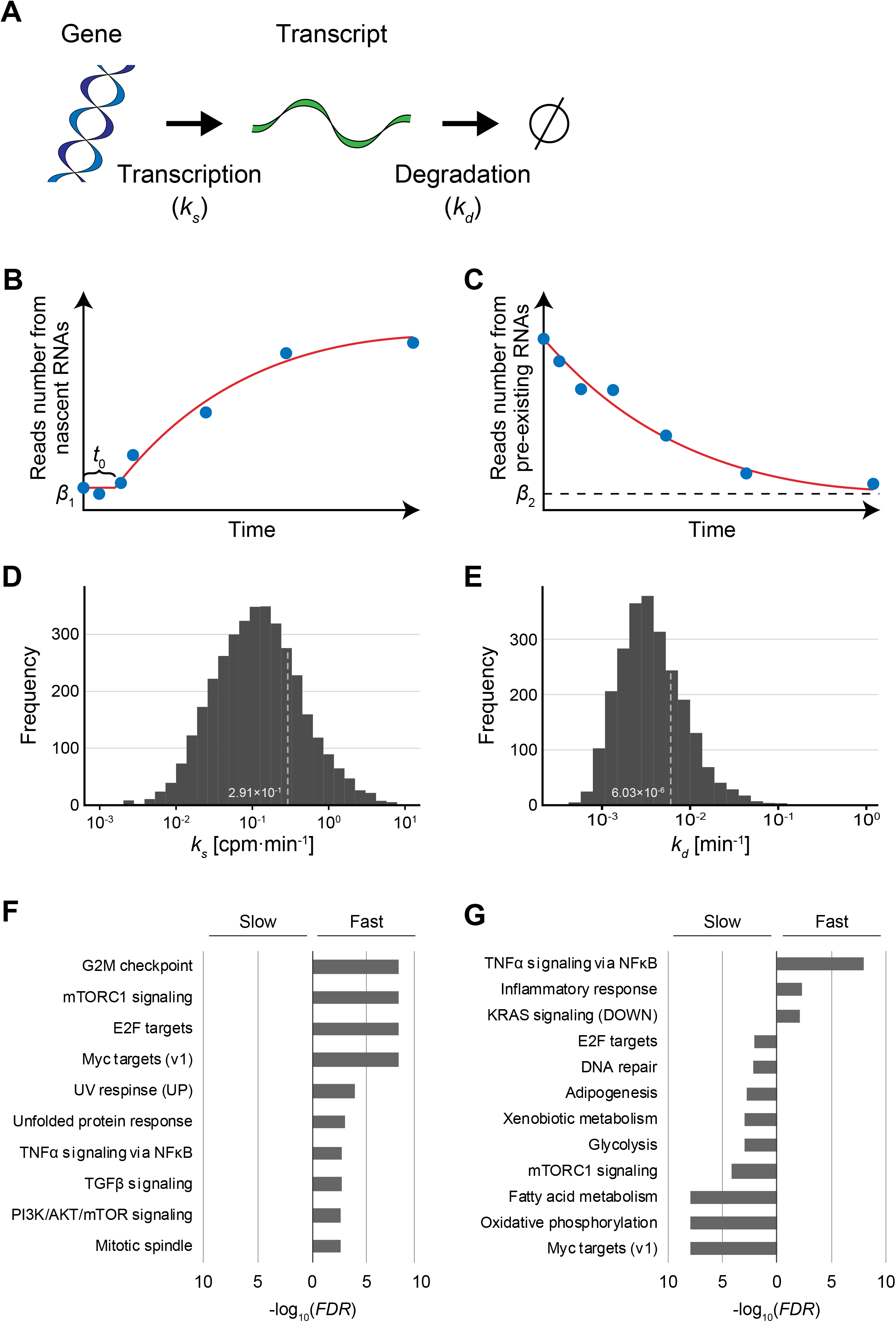
Estimation of transcription and degradation rates of individual genes. (**A**) Kinetics model of gene expression. In this model, a transcript was synthesized according to the transcription rate, *k_s_* [cpm·min^−1^], and degraded according to the degradation rate, *k_d_* [min^−1^]. (**B**) Transcription rates were estimated by fitting of the time series of the amount of nascent RNAs from individual genes to a logarithmically increasing curve (see STAR Methods). *β_1_* and *t*_0_ indicate the basal value of nascent RNAs and the time delay, respectively. (**C**) Degradation rates were estimated by fitting of the time series of the amount of pre-existing RNAs from individual genes to an exponentially decreasing curve (see STAR Methods). *β*_2_ indicates the basal value of pre-existing RNAs. (**D**and **E**) Distribution of estimated transcription rate (**D**) and degradation rate (**E**). (**F**and **G**) Gene Set Enrichment Analysis (GSEA) for genes with *k_s_* values (**F**) or with *k_d_* values (**G**).

Are these rates of transcription and degradation related to gene function? To approach this issue, we performed Gene Set Enrichment Analysis (GSEA) (Mootha *et al*., 2003; Subramanian *et al*., 2005) for the estimated transcription and degradation rates (Table S2). GSEA statistically tests the homogeneity of the *k_s_* values (transcription rates) and *k_d_* values (degradation rates) of genes related to specific biological terms: a uniform distribution of the *k_s_* or *k_d_* values of genes related to a term makes the *p*-value larger, while a biased distribution of these values makes it smaller. For the transcription rates, GSEA detected that the genes exhibiting only large *k_s_* values (fast transcription) are particularly related to several signaling pathways, such as mTORC1 signaling, TNFα signaling, TGFβ signaling, and PI3K/AKT/mTOR signaling (Fig. 2F). For the degradation rates, GSEA detected that the genes exhibiting only small *k_d_* values (slow degradation) are particularly related to several metabolic pathways, such as adipogenesis, xenobiotic metabolism, glycolysis, and fatty acid metabolism (Fig. 2G). TNFα signaling and mTORC1 signaling were related only to genes exhibiting large *k_d_* values (fast degradation) and those with small *k_d_* values (slow degradation), respectively (Fig. 2G). An integrated interpretation of these functional analyses indicates the following: The genes involved in the inflammatory response (TNFα signaling via NFκB, inflammatory response) are transcribed faster and their transcripts are degraded faster; the genes involved in cell growth and survival [E2F targets, Myc targets (v1), mTORC1 signaling] are transcribed faster but their transcripts are degraded slower (Fig. 2F and G). These results indicate that the rates of transcription and degradation are closely related to the biological function of the gene, and especially signaling factors tend to be transcribed faster and metabolic enzymes tend to be degraded slower.

### Comparison of transcription and degradation rates

To examine how the combination of transcription and degradation rates is related to the function and behavior of individual genes, we classified the 4,702 RNAs into four classes based on their transcription and degradation rates; fast transcription and fast degradation (Class I, 277 genes), fast transcription and slow degradation (Class II, 372 genes), slow transcription and fast degradation (Class III, 291 genes), and slow transcription and slow degradation (Class IV, 229 genes) (Table S3). To examine the relationship of the combination of transcription and degradation rates with the biological function of the genes, we performed functional enrichment analysis of the genes included in each class. This analysis of the genes using the DAVID tool (Huang, Sherman and Lempicki, 2009b, 2009a) provided several functional terms significantly enriched in the individual classes (Fig. 3A and B, Table S4). In Class I (fast transcription and fast degradation), the terms related to signaling pathways such as “serine/threonine-protein kinase” and those related to DNA repair were significantly enriched. In Class II (fast transcription and slow degradation), the terms were related to post-transcriptional regulation such as “mRNA splicing”, “mRNA processing”, and “RNA-binding”. The terms related to some signaling pathways such as NFκB signaling and Wnt signaling were also significantly enriched in Class II. The terms “alternative splicing” and “phosphoprotein” (indicating proteins to be phosphorylated) were significantly enriched in both Class I and Class II (classes of genes with fast transcription). The terms of transcriptional regulation including “zinc-finger” were significantly enriched in both Class I and Class III (classes of genes with fast degradation). The terms “acetylation” and “mitochondrion” were significantly enriched in both Class II and Class IV (classes of genes degraded slowly).

**Figure 3.**
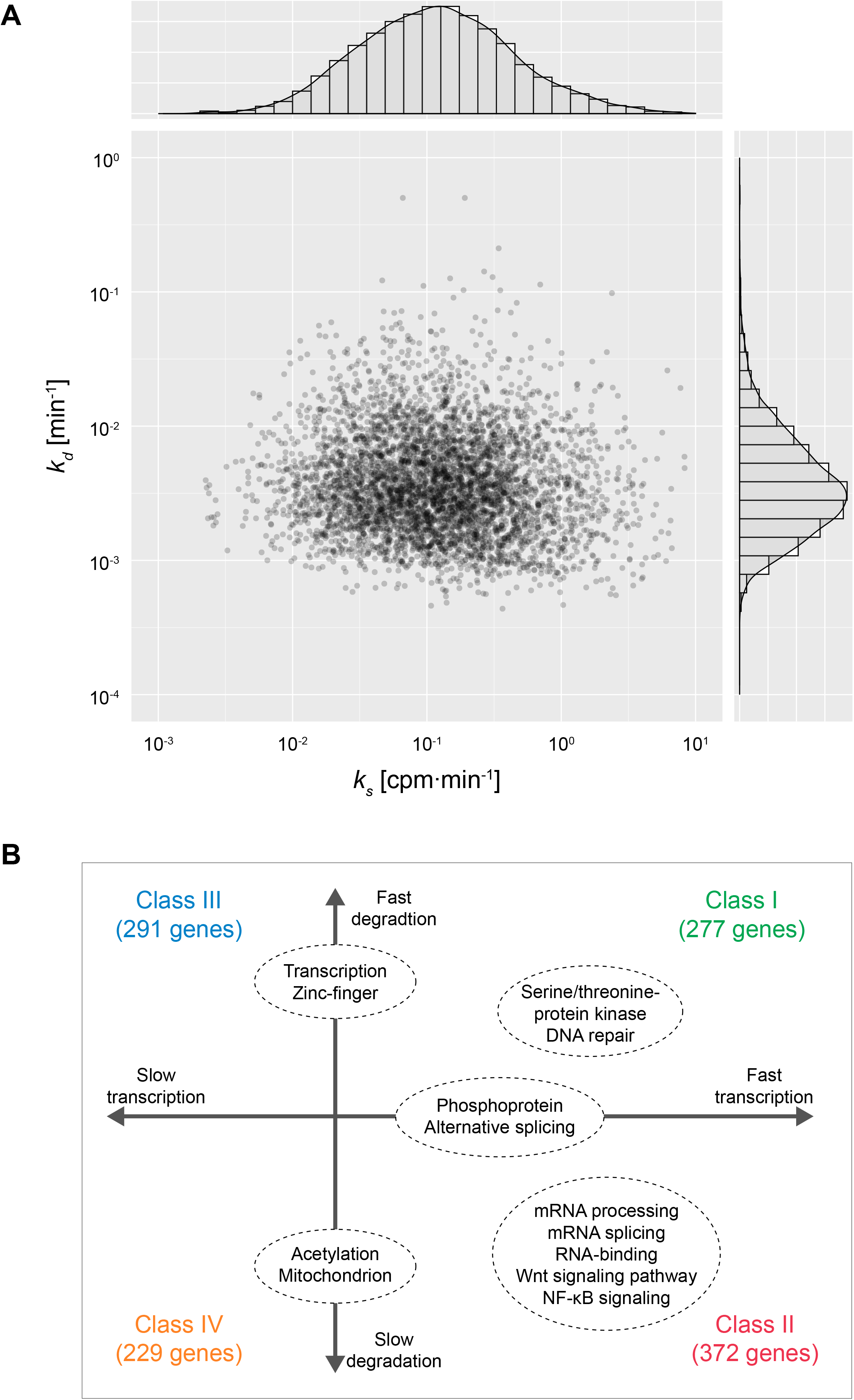
Joint distribution of transcription and degradation rates. (**A**) Joint distribution of transcription and degradation rates. Histograms of individual rates and a scatter plot are shown. X- and Y-axes indicate transcription rate (*k_s_*) and degradation rate (*k_d_*), respectively. (**B**) GO enrichment analysis of the genes classified using the transcription and degradation rates. Only representative terms significantly enriched for each class are shown.

In summary, functional enrichment analysis suggested the following: (i) The genes related to signaling are generally transcribed rapidly, which is consistent with the results of GSEA, but the rapidity of degradation varies depending on the signaling pathways. (ii) The genes related to post-transcriptional regulation such as splicing are transcribed rapidly and degraded slowly. (iii) The genes related to transcriptional regulation are degraded rapidly, while those encoding proteins to be acetylated are degraded slowly. These results show that genes have optimized transcription and degradation rates according to their functions and physiological roles.

### Combination of transcription and degradation rates determines expression level

Theoretically, the expression level of a gene in a steady state is determined by the ratio of its transcription and degradation rates (Hargrove and Schmidt, 1989). Thus, we examined how the transcription and degradation rates of individual genes affect the expression level using these estimated rates.

Generally, the expression rate of genes whose regulation is expressed as per the model in Fig. 2A is described as follows:

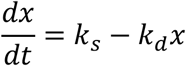

where *x*, *t*, *k*_*s*_, and *k*_*d*_ represent the expression level, time, transcription rate, and degradation rate of an RNA, respectively. Since the expression level does not change during a steady state,

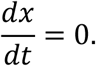

Thus, the expression level of the RNA in a steady state is described as:

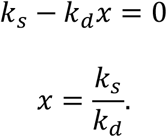

To verify the appropriateness of the theoretical prediction, we compared the experimentally estimated *k_s_* and *k_d_* values with the expression levels of individual genes estimated based on the sequencing of the Alkyl-RNAs. As expected, we observed strong positive correlation between the ratios of *k_s_* and *k_d_* values (*k_s_*/*k_d_*) and the expression levels (Pearson’s correlation of R^2^ = 0.85, *p*-value < 0.01) (Fig. 4A), suggesting that the combination of transcription and degradation rates determines the expression levels of individual genes. This indicates that not only transcriptional regulation but also the regulation of degradation is an important factor determining the level of gene expression.

**Figure 4.**
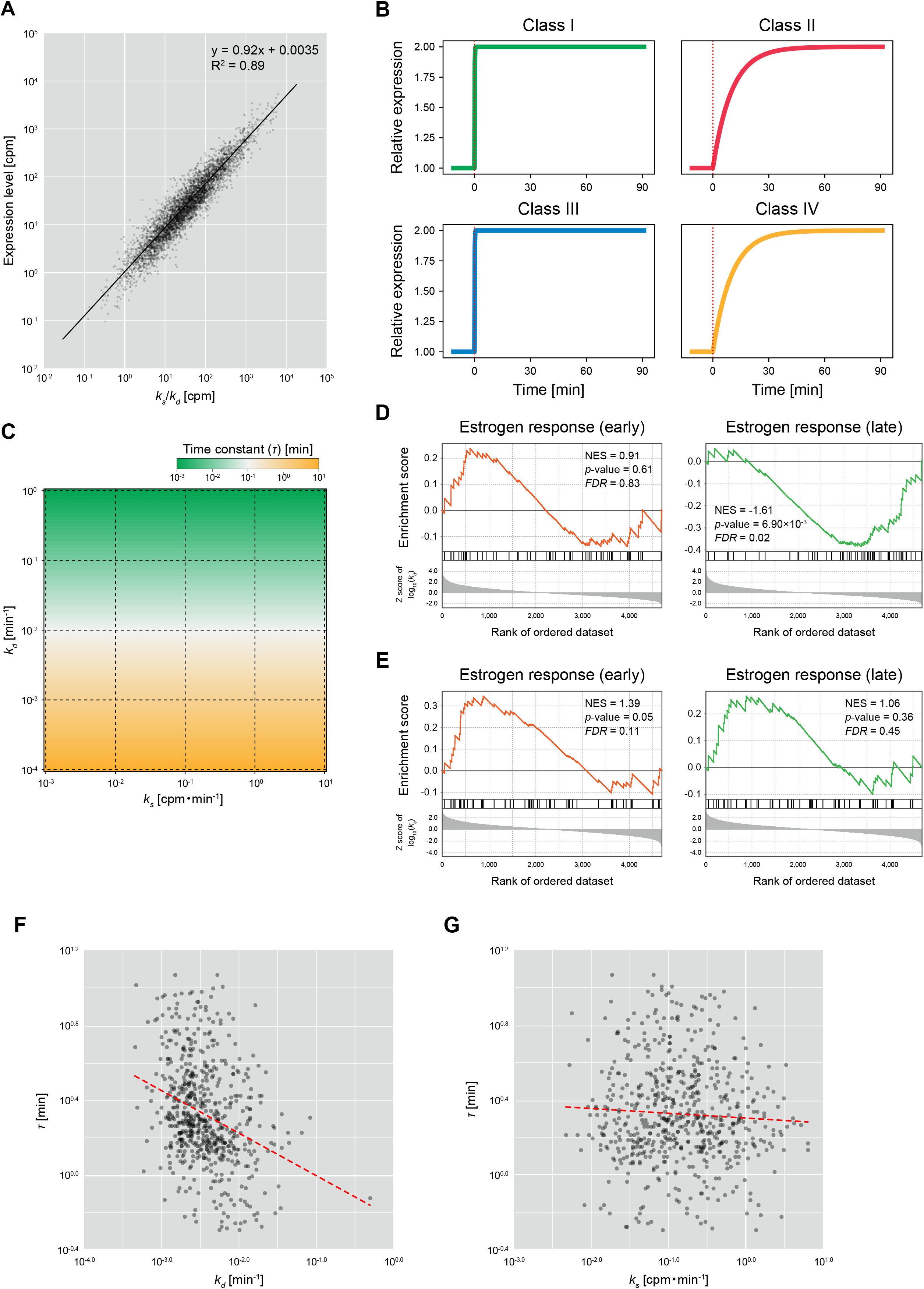
Effect of transcription and degradation rates on behavior of gene expression. (**A**) Comparison of ratios of transcription and degradation rates, with expression levels. X- and Y-axes indicate the ratio of transcription and degradation rates (*k_s_*/*k_d_*) and the expression level calculated based on the sequencing of Alkyl-RNAs (total RNA-seq reads with/without T > C conversion), respectively. Black solid line is the regression line based on log-transformed values. (**B**) Simulations of expression time series of genes imitating the characteristics of Class I (fast transcription and fast degradation), Class II (fast transcription and slow degradation), Class III (slow transcription and fast degradation), and Class IV (slow transcription and slow degradation). In each simulation, the transcription rates doubled at 0 min (red dotted line). (**C**) Time constants of expression of genes exhibiting various transcription and degradation rates. X- and Y-axes indicate transcription rate (*k_s_*) and degradation rate (*k_d_*). Colors indicate the time constants (*τ*). (**D**and **E**) GSEA of degradation rate (**D**) and transcription rate (**E**) showing enrichment of “estrogen response (early)” and “estrogen response (late).” NES, normalized enrichment score. (**F**) Comparison of degradation rates (*k_d_*) and time constants (*τ*). X- and Y-axes indicate the degradation rate and time constants, respectively. Red dashed line is a regression line based on log-transformed values. (**G**) Comparison of transcription rates (*k_s_*) and time constants (*τ*). X- and Y-axes indicate the transcription rate and time constants, respectively. Red dashed line is a regression line based on log-transformed values.

### Degradation rate is a key factor determining the rapidity of response

Finally, to examine how the transcription and degradation rates affect the behavior of individual RNAs (Koh, Porter and Batchelor, 2019), we simulated RNA expression dynamics in response to a change of transcriptional regulation using the mathematical model shown in Fig. 2A, and numerically analyzed the behavior of RNA expression. The dynamics of gene expression was simulated independently for four classes whose *k_s_* and *k_d_* values imitate the representative values of those in Classes I to IV. In this simulation, the transcription rate doubled at 0 min. The rapidity of expression was evaluated using a time constant (*τ*) defined as the time when the change of expression level reached 1−e^−1^ (≈ 63.2%) of the final value.

Theoretically, the time constant (*τ*) is represented as follows:

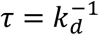

depending on not the transcription rate but only the degradation rate, indicating that the time constants are larger for the genes in the classes whose degradation is fast, such as Classes I and III, and they are smaller for the RNAs in the classes whose degradation is slow, such as Classes II and IV. In accordance with the theoretical prediction, the time constants simulated in Classes I and III (fast degradation) are smaller than those simulated in Classes II and IV (slow degradation) (Fig. 4B). Moreover, the time constants were determined depending on *k_d_*, not on *k_s_* (Fig. 4C). These results suggest that the degradation rate of each gene is a key factor determining not only their expression level but also their rapidity of differential expression caused by changes in the extracellular environment.

To confirm the above idea experimentally, we compared the estimated degradation rates with the time constants calculated based on previously published RNA-seq data. The GSEA for the degradation rates indicated that, while “estrogen response (early)” was significantly enriched for the large *k_d_* values (fast degradation), “estrogen response (late)” was significantly enriched for the small *k_d_* values (slow degradation) (*FDR* < 0.01) (Fig. 4D). In contrast, the GSEA for the transcription rates exhibited a uniform distribution of those terms (Fig. 4E). This indicates that the products of most genes related to late periods in estrogen response are degraded slowly, suggesting that the rapidity of expression of genes in response to estrogen is determined by their degradation rates. Since estrogen receptor is a nuclear receptor directly regulating transcriptional activity and its target genes might not be affected by the rapidity of signal transduction, the rapidity of expression of genes in response to estrogen is likely to reflect the degradation rates of the products of individual genes. Therefore, we calculated time constants from the time series of RNA-seq of human breast cancer-derived MCF-7 cells stimulated with estradiol (E2), a type of estrogen (Baran-Gale, Purvis and Sethupathy, 2016), and compared those with the estimated degradation rates. To avoid the effect of outliers of estimated expression levels, we redefined the time constant (*τ*) as the time when the change of expression level reaches half of the final value.

As expected, we observed a significant negative correlation between the degradation rates and time constants (Pearson’s correlation of *r* = −0.326, *p*-value = 1.65×10^−17^) (Fig. 4F), while the transcription rates were not significantly correlated (Pearson’s correlation of *r* = 0.033, *p*-value = 0.40) (Fig. 4G). This suggests that the rapidity of the response of gene expression to estrogen stimulation is at least partially determined in a manner dependent on the degradation rate. However, some genes exhibited extremely fast or slow time constants independent of their degradation rates, suggesting that the rapidity of expression of such genes affects secondary signal transduction and/or other signaling pathways.

## Discussion

Recent developments in procedures to unravel the kinetics of gene expression at the genome-wide level provide several insights into the regulation of gene expression. For instance, BRIC-seq (Tani *et al*., 2012; Imamachi *et al*., 2014) enables comprehensive clarification of the regulation of degradation through chase experiments of BrU-labeled RNAs. Moreover, SLAM-seq (Herzog *et al*., 2017) enables comprehensive clarification of transcriptional regulation through the quantification of 4sU-labeled RNAs by identifying IAA-induced T > C mutations. In this study, we developed “Dyrec-seq”, a system to simultaneously measure transcription and degradation rates of the comprehensive set of RNAs in HeLa cells, by chasing BrU- and 4sU-labeled RNAs. It provided simultaneous measurement of the transcription and degradation rates of 4,702 RNAs. Moreover, functional enrichment analysis of the genes classified by transcription and degradation rates demonstrated that these rates of genes related to various cellular functions are regulated in common, suggesting that the transcription and degradation rates of individual genes are closely associated with their biological functions. To examine how such regulation is associated with the biological roles of the genes, we constructed a mathematical model. This theoretical approach indicated that (i) the ratios of transcription rates to degradation rates of individual genes determine their expression levels, and (ii) the degradation rate is a key factor determining the rapidity of expression of each gene. Taking these results together, the following insights can be obtained (Fig. 5A and B): (i) Post-transcriptional factors including mainly RBPs, which are transcribed rapidly and degraded slowly, are expressed constitutively at extremely high levels, but respond slowly to extracellular stimuli. (ii) Transcription factors, which are degraded rapidly, exhibit various expression levels but respond to extracellular stimuli rapidly. (iii) The genes encoding phosphorylated proteins, including mainly signaling factors, which are transcribed rapidly, are generally expressed constitutively at extremely high levels, but their rapidity of response to extracellular stimuli vary according to the related signaling pathways. (iv) The genes encoding proteins under post-translational regulation, such as acetylation, which are degraded slowly and do not respond rapidly to extracellular stimuli in terms of the gene expression level, might mainly be regulated post-translationally. These insights explain at least in part how the differences of expression levels are regulated, and why the responses to extracellular stimuli differ according to the function of the genes. Simultaneous measurement of transcription and degradation rates using Dyrec-seq provides an integrated understanding of the functions of genes and their behavior of expression.

**Figure 5.**
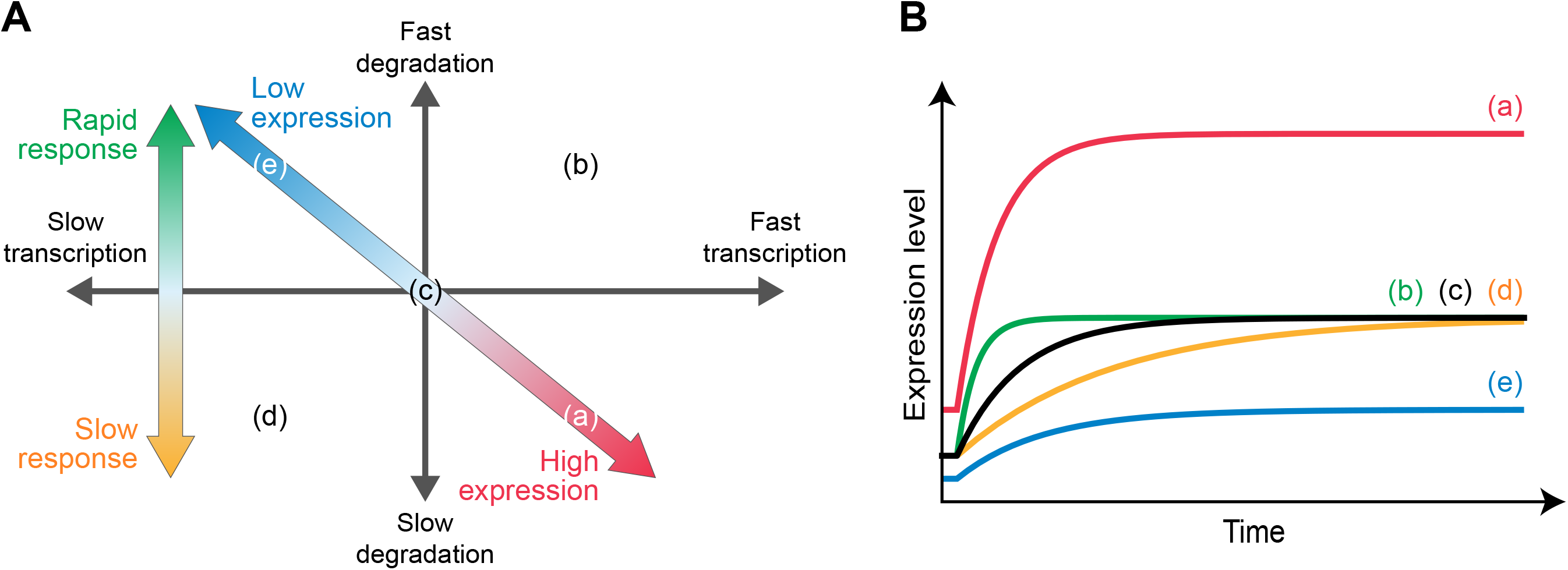
Effect of transcription and degradation rates on behavior of gene expression. (**A**) The combination of the rates of transcription and degradation regulates the temporal expression profiles. The ratio of the transcription and degradation rates determines the expression level (diagonal arrow), and the degradation rate affects the rapidity of response of gene expression (vertical arrow). (**B**) Schematic diagram of temporal expression profiles. The temporal expression profiles correspond to the points in **Figure 5A**.

To measure the transcription and degradation rates simultaneously, we labeled the intracellular RNAs using two uridine analogs, BrU and 4sU. Since it is known that uridines are particularly abundant in the 3′ untranslated region (3′UTR) in RNAs (Shaw and Kamen, 1986; Chen and Shyu, 1995; Barreau, Paillard and Osborne, 2005; Vlasova and Bohjanen, 2008; Vlasova *et al*., 2008; Gruber *et al*., 2011), we used QuantSeq, which specifically detects 3′UTR using oligo(dT) primer (Moll *et al*., 2014), to ensure efficient detection. Note that the transcription rates that we measured in this study include inherent transcription rates and processing rates. The efficiency of incorporation of nucleotide analogs and the effects of cellular functions might be influenced by the cell type and conditions. Thus, there is a need to verify effects specific to cell types sufficiently to identify labeling conditions that do not influence the gene expression of cells. Moreover, the ability to estimate the transcription and degradation rates depends on the inherent kinetics of gene expression and library sequencing depth. Therefore, consideration of these parameters is needed to design experiments effectively.

Further extension of Dyrec-seq with other kinds of nucleotide analogs would enable the evaluation of additional types of regulation such as splicing and processing (Windhager *et al*., 2012; Barrass *et al*., 2015). Moreover, this procedure can be used for understanding the temporal contributions of multiple types of regulation such as transcription and degradation during transient or continuous changes of gene expression. Recent studies suggested that the regulation of both transcription and degradation can affect the dynamics of gene expression (Alonso, 2012). For example, upon regulation of the gene expression profile in the long term, such as occurs in cell differentiation, the contribution of mechanisms regulating transcription and degradation might change depending on the stage of differentiation.

In conclusion, we propose the concept of “RNA dynamics recording” to encode the dynamics of RNAs onto the RNAs themselves by using multiple ribonucleoside analogs. This RNA dynamics recording enables clarification of the temporal contributions of multiple forms of regulation independently. Therefore, this approach should provide substantial insights into how gene expression is regulated during differentiation or by extracellular stimuli such as hormones and cytokines.

## Acknowledgments

We thank our laboratory members for critically reading this manuscript and for their technical assistance with the experiments. We also thank Kouta Watanabe, Kiyomi Imamura, Terumi Horiuchi, and Yuta Kuze for performing NGS experiments and analysis. The computational analysis of this work was performed on the National Institute of Genetics (NIG) supercomputer system at Research Organization of Information and Systems (ROIS). This manuscript was edited by Edanz (www.edanzediting.co.jp). This work was supported by the Japan Society for the Promotion of Science (JSPS) KAKENHI (Grant Number: 17KK0163, 18H02570, 18KT0016, and 16H06279). K.K. received funding from JSPS KAKENHI (Grant Number: 19K16635) and Takeda Life Science Foundation. T.Y. received the JSPS Research Fellowship for Young Scientists and is supported by a Grant-in-Aid for JSPS Fellows (17J11266). M.S. was supported by JSPS KAKENHI (Grant Number 16H06279 (PAGS) and 19K16108). Y.S. was supported by JSPS KAKENHI (Grant Number 16H06279 (PAGS)).

## Author Contributions

K.K. and N.A. conceived the project. H.W., H.H., and N.A. designed and performed the experiments. K.K., T.Y., and K.T. analyzed the data. M.S. and Y.S. performed the RNA-seq experiments. K.K, T.Y., K.T., and N.A. wrote the manuscript.

## STAR Methods

### Metabolic labeling of endogenous RNAs and RNA isolation

HeLa cells (female, RRID: CVCL_0030) were seeded at a density of 6×10^5^ cells per 6-cm dish (Thermo Fisher Scientific) and cultured in Dulbecco’s Modified Eagle’s Medium (DMEM) supplemented with 10% fetal bovine serum (FBS) and 1% antibiotic–antimycotic at 37ºC. After 12 h, the medium was replaced with fresh medium containing 150 μM BrU and then the cells were pre-cultured for 12 h to label pre-existing RNAs. After the pre-culture, the cells were washed twice with the medium, and then the medium was replaced with one containing 200 μM 4sU to label nascent RNAs. After incubating cells in the 4sU-containing medium, the cells were washed twice with PBS and then total RNA was isolated using RNAiso Plus (TAKARA).

### Sample preparation

To quantify 4sU- and BrU-labeled RNAs individually, the isolated total RNAs were divided into two samples, as follows.

#### RNA samples to quantify 4sU-labeled RNAs

From the isolated total RNAs, 5 μM RNA was alkylated using SLAMseq Kinetics Kit (Lexogen), in accordance with the manufacturer’s instructions. Hereafter, the alkylated RNA samples are referred to as “Alkyl-RNAs”. The quality of Alkyl-RNAs was assessed using Agilent RNA Nano 6000 Kit (Agilent) on the Agilent BioAnalyzer 2100 (Agilent).

#### RNA samples to quantify BrU-labeled RNAs

To normalize the quantity of BrU-labeled RNAs among the time points during pre-analysis, 1.0 ng of BrU-labeled luciferase RNA was added to 10 μg of extracted total RNA as a spike-in control to serve as an internal standard. The RNA mixtures were diluted to a final volume of 100 μl with TE buffer (a final concentration of 10 mM Tris–HCl at pH 7.0 and 1.0 mM EDTA). BrU-labeled RNAs were immunoprecipitated using anti-BrdU antibody-conjugated beads, in accordance with previous reports (Tani *et al*., 2012; Imamachi *et al*., 2014). Briefly, the BrU-labeled RNAs were added to the antibody-conjugated protein G agarose suspended with 100 μl of ice-cold PBS containing 0.1% Bovine Serum Albumin (BSA), 1% Triton X100, 100 U of RNasin plus RNase inhibitor (Promega), and 5 mg/mL heparin. BrU-labeled RNAs were isolated using ISOGEN LS (NIPPON GENE), in accordance with the manufacturer’s instructions. Since the amount of purified BrU-labeled RNAs was low, 60 μg of glycogen was added to the mixture during the precipitation of RNA. Hereafter, the immunoprecipitated RNA samples are referred to as “IP-RNAs.” The quality of IP-RNAs was assessed using Agilent RNA Nano 6000 Kit (Agilent) on the Agilent BioAnalyzer 2100 (Agilent).

### Sequencing

The 3′ ends of RNAs within the Alkyl- and IP-RNAs were sequenced using the QuantSeq 3′ mRNA-Seq Library Prep Kit (Lexogen), in accordance with the manufacturer’s instructions. Briefly, 5.0 ng from each Alkyl- or IP-RNA sample was used for reverse transcription. The 4sU incorporated into the Alkyl-RNAs paired with guanines instead of adenines during the reverse transcription. The RNA was subsequently removed, and second-strand synthesis was initiated by a random primer, containing Illumina-compatible linker sequences and appropriate in-line barcodes, followed by magnetic bead-based purification. The resulting library was amplified using PCR with 12 cycles for Alkyl-RNAs and 18 cycles for IP-RNAs, and then purified using AMPure XP. Library quality and quantity were assessed on a BioAnalyzer using DNA High Sensitivity Kit reagents (Agilent Technologies). Standard Illumina protocols were used to generate 100-base pair-end read libraries that were sequenced on the HiSeq3000 platform (Illumina).

### Quantification of 4sU- and BrU-labeled RNAs

The nascent and pre-existing RNAs were quantified based on the sequence data obtained from the Alkyl- and IP-RNAs for each time point by using SlamDunk v0.3.3, a pipeline for SLAM seq data analysis, with the default parameters (Herzog *et al*., 2017). Since QuantSeq targets mainly the 3′ end of individual RNAs in a poly(A) tail-dependent manner, the sequence data were aligned on genome-wide 3′UTR sequences generated based on the human genome sequence and annotation data (GRCh38) obtained from the Ensembl database (release 92). Twelve bases from the 5′ end were trimmed as adaptor-clipped reads, and then four or more subsequent adenines from the 3′ end were regarded as the remaining poly(A) tail and removed. Multiply mapped reads were allowed up to 100 of regions. In VarScan2.4.1 (Koboldt *et al*., 2012) included in the SlamDunk tool, an SNP was called in the case of a mismatch exceeding a variant fraction of 0.8 and a coverage cut-off of 10-fold. Through these filters, we calculated the total number of reads and the number of those including non-SNP T > C conversions aligned on the 3′UTR of individual genes. Since 4sU pairs with guanine (G) during reverse transcription, the 4sU-labeled reads were identified as those including T > C conversion. Since one read is generated from one RNA, the total number of reads and the number of those including T > C conversions corresponds to the numbers of total RNAs and 4sU-labeled RNAs, respectively. Therefore, the numbers of reads including T > C conversion from Alkyl-RNAs and those of reads from IP-RNAs were counted as the numbers of 4sU- and BrU-labeled RNAs, respectively, and normalized to counts per million mapped reads (CPM).

### Identification of RNAs in a steady state

Although we did not stimulate the cells in this study and the genes should have been in a steady state, we observed that some of the genes exhibited differential expression. Since the expression of a gene in a steady state changes dependent on only white noise, the sum of angles formed by the lines connecting each time point is relatively small, while the sum of angles in a differentially expressed gene with a constant trend is relatively large (Fig. 1F). Therefore, we designed a filter to exclude the differentially expressed genes with constant trends. First, we calculated the sum of the angles formed by the lines connecting certain time points and neighboring ones, within a time series of total read numbers for each gene calculated from Alkyl-RNAs. Second, we generated an empirical distribution of the sum of angles by rearranging all time points randomly. Finally, we calculated the probability that the sum of angles in the empirical distribution was larger than the actual value (empirical *p*-value). We calculated FDR from the empirical *p*-values using Storey’s procedure (Storey, Taylor and Siegmund, 2004). Among the genes expressed at all time points, those with *FDR* values greater than 0.01 were identified as genes in a steady state.

### Correction of nascent and pre-existing RNA numbers

Since the 4sU added to the medium labels only part of nascent RNAs due to the incorporation efficiency, we estimated the incorporation efficiency based on the time series of total 4sU-labeled RNAs. In a steady state, constant copies of 4sU-labeled RNAs are generated and their level decreases at a constant rate per unit time. Therefore, the time series of 4sU-labeled RNAs in count per million mapped reads (CPM) obeys the following:

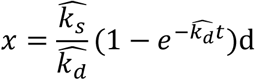

where *x,* 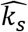, and 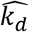 indicate the number of 4sU-labeled RNAs in CPM, the transcription rate, and the degradation rate of the total 4sU-labeled RNAs, respectively. In addition, this time series asymptotically approaches the ratio of 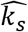 to 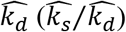. Since the number of nascent RNAs in CPM can approximate 10^6^, the incorporation efficiency (*η*) can be presented as:

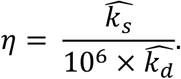

Therefore, the numbers of nascent RNAs of total and individual genes are calculated by dividing the number of 4sU-labeled RNAs in CPM by the estimated incorporation efficiency.

Moreover, we estimated the number of pre-existing RNAs at each time point. Since the number of intracellular RNAs in a steady state can be regarded as constant, the total number of pre-existing RNAs in CPM can be determined by subtracting the total number of nascent RNAs in CPM from 1 million. Therefore, the number of pre-existing RNAs of individual genes was calculated by correcting the number of BrU-labeled RNAs of individual genes in CPM by the total number of pre-existing RNAs.

### Estimation of transcription and degradation rates

The numbers of nascent and pre-existing RNAs increase and decrease, respectively, in a time-dependent manner, according to the transcription and degradation rates. Therefore, we estimated the transcription and degradation rates simultaneously by fitting the time series of the estimated nascent and pre-existing RNA levels to:

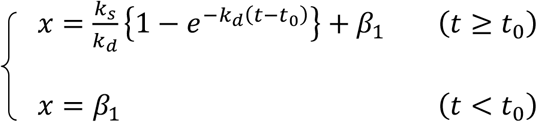

and

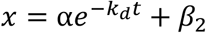

where *k_s_*, *k_d_*, *α*, *t*_0_, *β_1_*, and *β_2_* indicate the transcription rate, degradation rate, scaling factor, time delay, basal value for nascent RNAs, and basal value for pre-existing RNAs of individual genes, respectively. The scaling factor is incorporated as the number of pre-existing RNAs at 0 min. The time delay is incorporated as the time when the nascent RNAs begin to be synthesized after 4sU addition. The basal value for nascent RNAs is incorporated as the number of RNAs including inherent T > C mutant. The basal value for pre-existing RNAs is incorporated as the background of immunoprecipitation of BrU-labeled RNAs.

The fittings were performed with the combination of an evolutionary algorithm (genetic algorithm) and hill climbing (L-BFGS-B algorithm) with evaluation by the least squares method in Python2.7. The genetic algorithm was implemented using the *DEAP* library with a generation number of 200, population number of 50, crossover probability of 0.5, and mutation probability of 0.2. The boundaries of parameters are shown in Table 1. The L-BFGS-B algorithm was implemented using the *minimize* module in the *SciPy* package, where the parameters estimated by the genetic algorithm are given as initial parameters. The fitness in each gene was evaluated as the correlation of actual nascent and pre-existing RNA levels with estimated values. The probability of the null hypothesis that a population correlation coefficient is equivalent to 0 was calculated for each gene using the *OLS* module in the *StatsModels* package, and the transcription and degradation rates of the genes whose *FDR* as determined by Storey’s procedure (Storey, Taylor and Siegmund, 2004) was less than 10^−5^ were extracted.

**Table 1.**
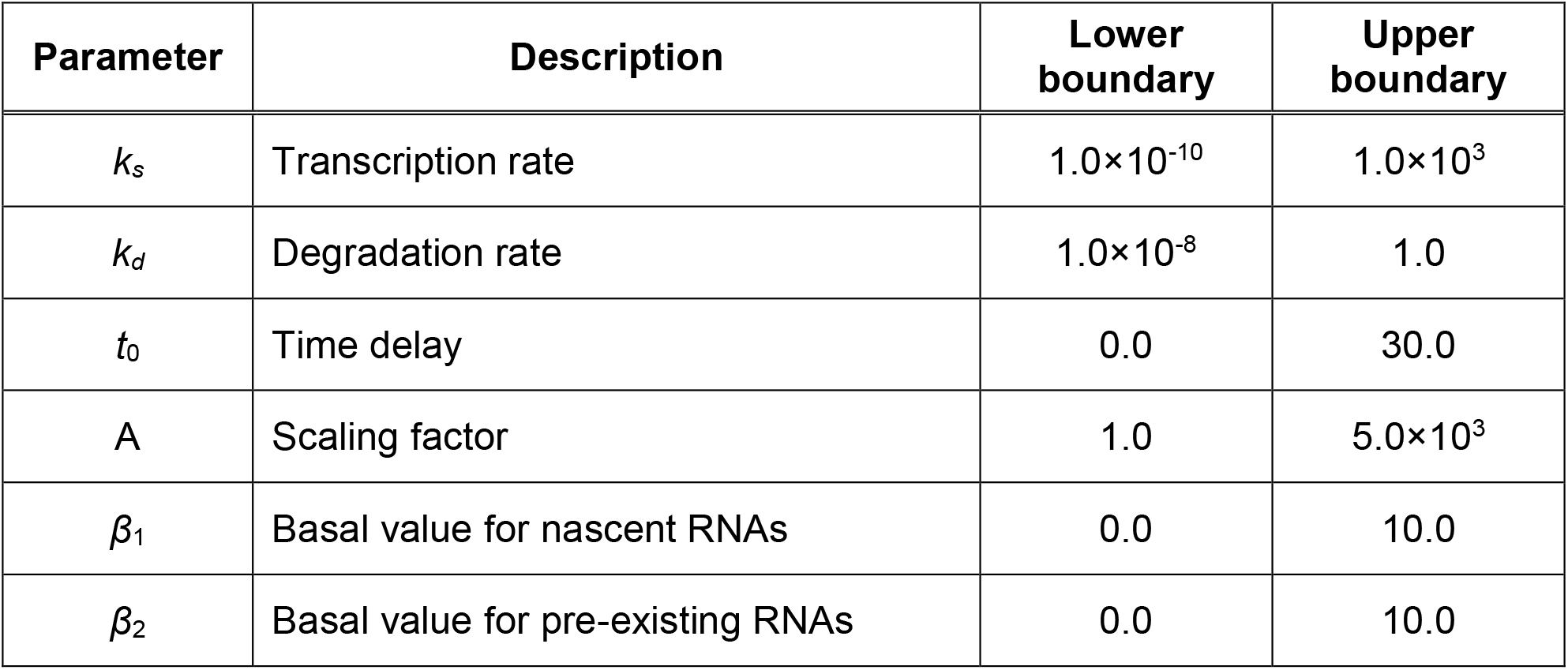
Boundaries of parameters.

| Parameter | Description | Lower boundary | Upper boundary |
| --- | --- | --- | --- |
| $k_s$ | Transcription rate | $1.0 \times 10^{-10}$ | $1.0 \times 10^3$ |
| $k_d$ | Degradation rate | $1.0 \times 10^{-8}$ | 1.0 |
| $t_0$ | Time delay | 0.0 | 30.0 |
| A | Scaling factor | 1.0 | $5.0 \times 10^3$ |
| $\beta_1$ | Basal value for nascent RNAs | 0.0 | 10.0 |
| $\beta_2$ | Basal value for pre-existing RNAs | 0.0 | 10.0 |

### Gene Set Enrichment Analysis (GSEA)

The functions of genes and their associations with the estimated transcription and degradation rates were independently examined using Gene Set Enrichment Analysis (GSEA) 4.0.3. We set 1 as the control value for individual genes, and sorted the genes by the ratio of the *k_s_* or *k_d_* values to the control values (i.e., the original *k_s_* or *k_d_* values). The enrichment scores (ESs) for the terms included in GSEA hallmark were calculated using the default parameters, and empirical *p*-values were calculated by comparing those with the distributions of ES values from 10,000 randomized gene sets. The empirical *p*-values were corrected as FDR. The terms with FDRs of less than 0.05 were considered as significantly enriched.

### Classification of genes according to transcription and degradation rates

The genes were classified according to the associated transcription and degradation rates. To avoid the effects of estimation error, first and third quartiles were adopted as thresholds of the classification. The first and third quartiles were calculated for the transcription rate (*k_s_*) and degradation rate (*k_d_*) independently: *Q_s1_*, first quartile of *k_s_* values; *Q_s3_*, third quartile of *k_s_* values; *Q_d1_*, first quartile of *k_d_* values; and *Q_d3_*, third quartile of *k_d_* values. Using these thresholds, we classified the genes into four classes: Class I, fast transcription (*k_s_* > *Q_s3_*) and fast degradation (*k_d_* > *Q_d3_*); Class II, fast transcription and slow degradation (*k_d_* < *Q_d1_*); Class III, slow transcription (*k_s_* < *Q_s1_*) and fast degradation; and Class IV, slow transcription and slow degradation.

### Functional enrichment analysis

The biological functions of the genes classified by the transcription and degradation rates were statistically determined using the DAVID tool v6.8 (https://david.ncifcrf.gov/) (Huang, Sherman and Lempicki, 2009b, 2009a), by examining the Gene Ontology (GO) categories of biological process (GOTERM_BP_DIRECT), cellular component (GOTERM_CC_DIRECT), and molecular function (GOTERM_MF_DIRECT), as well as UniProt Keywords (UP_KEYWORDS). The *p*-values of enrichment were calculated by modified Fisher’s exact test (Fisher, 1922; Huang, Sherman and Lempicki, 2009b, 2009a). The whole human genome (*Homo sapiens*) was used as a background (default). The biological functions whose FDR values were less than 0.01 were identified as those that were significantly enriched.

### Estimation of time constants from published database

The time constants were calculated as a criterion of the rapidity of the response of gene expression. To calculate the time constants, a pre-quantified data set of RNA-seq of human breast cancer-derived MCF-7 cells stimulated with E2 (Baran-Gale, Purvis and Sethupathy, 2016) was downloaded from Gene Expression Omnibus (GEO) (https://www.ncbi.nlm.nih.gov/geo/query/acc.cgi?acc=GSE78169). This pre-quantified RNA-seq data set consists of the expression levels at 0, 1, 2, 3, 4, 5, 6, 8, 12, and 24 h after E2 stimulation with triplicate data for individual time points. First, the significance of the changes of the expression levels at each time point after stimulation relative to those without stimulation (0 h) was tested by Welch’s *t*-test using the *ttest_ind* function of the *stats* module in *Scipy* library (Virtanen *et al*., 2020). The genes whose expression levels were significantly changed in at least one time point were extracted as differentially expressed genes (DEGs). For each DEG, the time constant (*τ*), which is the time when the change of expression level reaches half of the final expression level, was calculated. The time constants whose identifiers were replaced with Ensembl Gene ID from NCBI Gene ID based on *db2db* on the bioDBnet web service (https://biodbnet-abcc.ncifcrf.gov/) (Mudunuri *et al*., 2009) were compared with the estimated transcription and degradation rates.

### Data and Code Availability

The BridgePy tool for simultaneous estimation of transcription and degradation rates is available from GitHub (https://github.com/kntr-kwt/BridgePy). The high-throughput sequencing data of Alkyl- and IP-RNAs have been deposited in the DDBJ database (https://www.ddbj.nig.ac.jp/index-e.html) as PRJDB8372.

## Supplemental Information titles and legends

Table S1: Estimated transcription and degradation rates

Table S2: Results of Gene Set Enrichment Analysis (GSEA)

Table S3: Classes of genes according to transcription and degradation rates

Table S4: Results of Gene Ontology (GO) enrichment analysis

